# Laminar profile of task-related plasticity in ferret primary auditory cortex

**DOI:** 10.1101/354910

**Authors:** Nikolas A. Francis, Diego Elgueda, Bernhard Englitz, Jonathan B. Fritz, Shihab A. Shamma

**Author notes:** Co-first authorship (Nikolas A. Francis, Diego Elgueda). Corresponding author: Nikolas A. Francis, Institute for Systems Research 2202 A.V. Williams Building University of Maryland College Park, MD 20742 USA.

## Abstract

Rapid task-related plasticity is a neural correlate of selective attention in primary auditory cortex (A1). Top-down feedback from higher-order cortex may drive task-related plasticity in A1, characterized by enhanced neural representation of behaviorally meaningful sounds during auditory task performance. Since intracortical connectivity is greater within A1 layers 2/3 (L2/3) than in layers 4-6 (L4-6), we hypothesized that enhanced representation of behaviorally meaningful sounds might be greater in A1 L2/3 than L4-6. To test this hypothesis and study the laminar profile of task-related plasticity, we trained 2 ferrets to detect pure tones while we recorded laminar activity across a 1.8 mm depth in A1. In each experiment, we analyzed current-source densities (CSDs), high-gamma local field potentials (LFPs), and multi-unit spiking in response to identical acoustic stimuli during both passive listening and active task performance. We found that neural responses to auditory targets were enhanced during task performance, and target enhancement was greater in L2/3 than in L4-6. Spectrotemporal receptive fields (STRFs) computed from CSDs, high-gamma LFPs, and multi-unit spiking showed similar increases in auditory target selectivity, also greatest in L2/3. Our results suggest that activity within intracortical networks plays a key role in shaping the underlying neural mechanisms of selective attention.

## Introduction

Selective attention is believed to facilitate auditory task performance by enhancing neural representation of behaviorally meaningful sounds^1–7^. Task-related plasticity is a neural correlate of selective attention that is characterized by transient changes in both the gain of neuronal responses to auditory targets, and the shape of spectrotemporal receptive fields (STRFs)^4,6,7^. During pure-tone detection and discrimination tasks, individual neurons become more responsive to auditory targets, while STRFs become more selective for the auditory target frequency^4,6,7^.

Converging lines of evidence from both anatomical and neurophysiological studies suggest that task-related plasticity in A1 may be greater in cortical layer 2/3 (L2/3) than layers 4-6 (L4-6), due to intracortical network activity within L2/3 that is believed to mediate top-down control of sensory processing^8–13^. The L2/3 intracortical network may provide a pathway for prefrontal cortex to bias A1 responsiveness in favor of behaviorally meaningful sounds^14–16^. However, the laminar profile of task-related plasticity in A1 remains unclear since few studies have recorded simultaneously across layers during auditory task performance^17,18^. In humans, behavioral detection of a frequency modulation sweep sharpened frequency tuning in superficial cortical layers more than in middle-deep layers^17^. In monkeys, current source density (CSD) analysis of local field potentials (LFPs) from A1 revealed that intermodal attention-related suppressive effects predominated the responses of superficial cortical layers, yet response enhancement was dominant in middle-deep layers^18^. Long-lasting effects of auditory training on A1 responses to sounds in anesthetized rats include an enhancement of CSD current sinks in L2/3, but not in layer 4^19^. It is believed that CSD amplitudes measure transmembrane currents related to neuronal spiking^20–22^, and that high-frequency LFPs (i.e. “high-gamma” LFPs >80 Hz) measure synchronous spiking from many neurons^20^. Here, we hypothesized that task-related plasticity might be (1) greater in superficial L2/3 than in middle-deep L4-6, and (2) similar for multi-unit spiking, high-gamma LFPs, and CSDs.

## Results

### Target response enhancement was greatest in L2/3

We studied the laminar profile of rapid task-related plasticity by recording from a 24 channel linear electrode array (Plexon U-probe) inserted through the dura, orthogonal to the surface of A1, in two ferrets that were performing an auditory detection task (Fig. 1a). During task performance trials, the animal heard a sequence of reference noises followed by a pure-tone target (Fig. 1b). Upon detecting the target, the animal was trained to stop licking a waterspout to avoid a mild shock^4^. Neural responses to the same sounds used during the task were also measured while the animal was in a passive, quiescent state, to provide a within-animal passive control condition for neural activity. Wide-band recordings from the 24 channel linear array allowed us to analyze multi-unit spiking, high-gamma LFP magnitudes, and CSD amplitudes across a 1.8 mm depth that included layers 2/3-6^12,23^(see Methods and Fig. 1c). All statistical comparisons were done using a bootstrap t-test.

**Figure 1.**
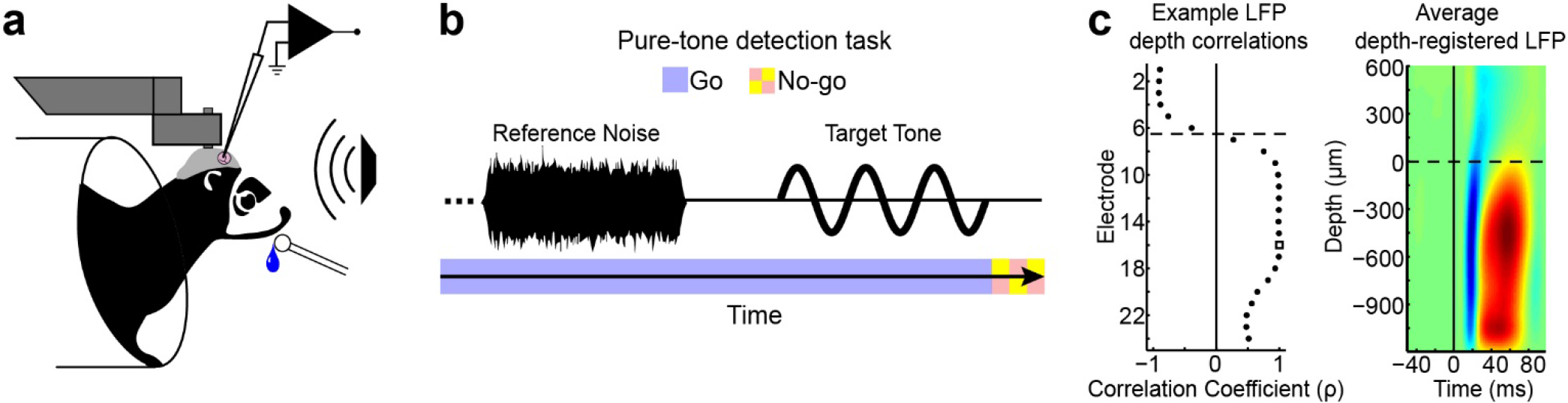
Awake-behaving experimental paradigm and electrode depth registration. **a.** Head-fixed preparation. Ferrets were implanted with a metal post that was attached to the skull and held fixed during awake-behaving neurophysiological experiments. The ferret performed the task while we recorded from primary auditory cortex (A1) using a 24 channel linear electrode array (Plexon U-probe). **b.** Pure-tone detection task. Two ferrets were trained to do a conditioned avoidance Go/No-Go pure-tone detection task. In each trial of the task, the animal heard a sequence of reference noises followed by a pure-tone target. Reference noises were “Go” signals, during which the animal was free to lick a waterspout. Upon detecting the target (the “No-Go” signal), the animal stopped licking the water spout to avoid a mild shock. The target frequency was different for each experiment. **c.** Electrode depth-registration. The left panel shows an example of how the layer 2/3 (L2/3) vs. layers 4-6 (L4-6) border (dashed line) was computed for a single penetration of the 24 channel linear electrode array in A1. Local field potential (LFP) responses to 100 ms tones were used to find a common marker of depth across penetrations (i.e., for depth registration). Registration began by first identifying the electrode with the shortest LFP response latency (Eτ, white square), then finding the LFP waveform correlation coefficients (ρ) between Eτ and all other electrodes in the same penetration. The border between the first neighboring electrode pair with positive and negative correlation coefficients defined the L2/3 vs. L4-6 border^12,22,23,38–40^. Laminar profiles were averaged across penetrations after first aligning to the border. The right panel shows the average depth-registered LFP laminar profile in response to 100 ms tones.

During task performance (average target detection d’=1.3, std=0.74), we found that neural responses to targets were enhanced relative to reference responses, and target enhancement was greater in superficial L2/3 than middle-deep L4-6. Figure 2 shows laminar profiles of the average responses to target and reference sounds, during passive and behavior conditions, for multi-unit spiking (top row), high-gamma LFPs (middle row), and CSDs (bottom row). Both target and reference sounds evoked responses across layers 2/3-6, during both passive and behavior conditions (Fig. 2a, d, and g). To quantify task-related plasticity (P) from neural responses (P_Resp_), we first computed the ratio of response amplitudes during behavior vs. passive trials, separately for target (Tar_Bhv/Pass_) and reference (Ref_Bhv/Pass_) sounds. Then we took the difference between target and reference ratios (P_Resp_ = Tar_Bhv/Pass_ - Ref_Bhv/Pass_; Fig. 2b,e,h). Positive values of P_Resp_ (red in figure 2b,e,h) indicate relative target response enhancement during auditory task performance. We found that target enhancement was the predominant effect for multi-unit spiking (Fig. 2b,c), high-gamma LFPs (Fig. 2e,f), and CSDs (Fig. 2h,i). Furthermore, target enhancement was greater in L2/3 electrodes for most recordings (see cumulative distribution functions in Fig. 2c,f,i). The average target enhancement values for L2/3 vs. L4-6 were: 0.48 vs. 0.09 (multi-unit spiking, p<0.001); 0.52 vs. 0.19 (high-gamma LFP, p<0.001); and 0.23 vs. 0.18 (CSD, p<0.05). The 9% average target enhancement we measured in multi-unit spiking from L4-6 agrees with previous measurements of multi-unit responses in A1^5^.

**Figure 2.**
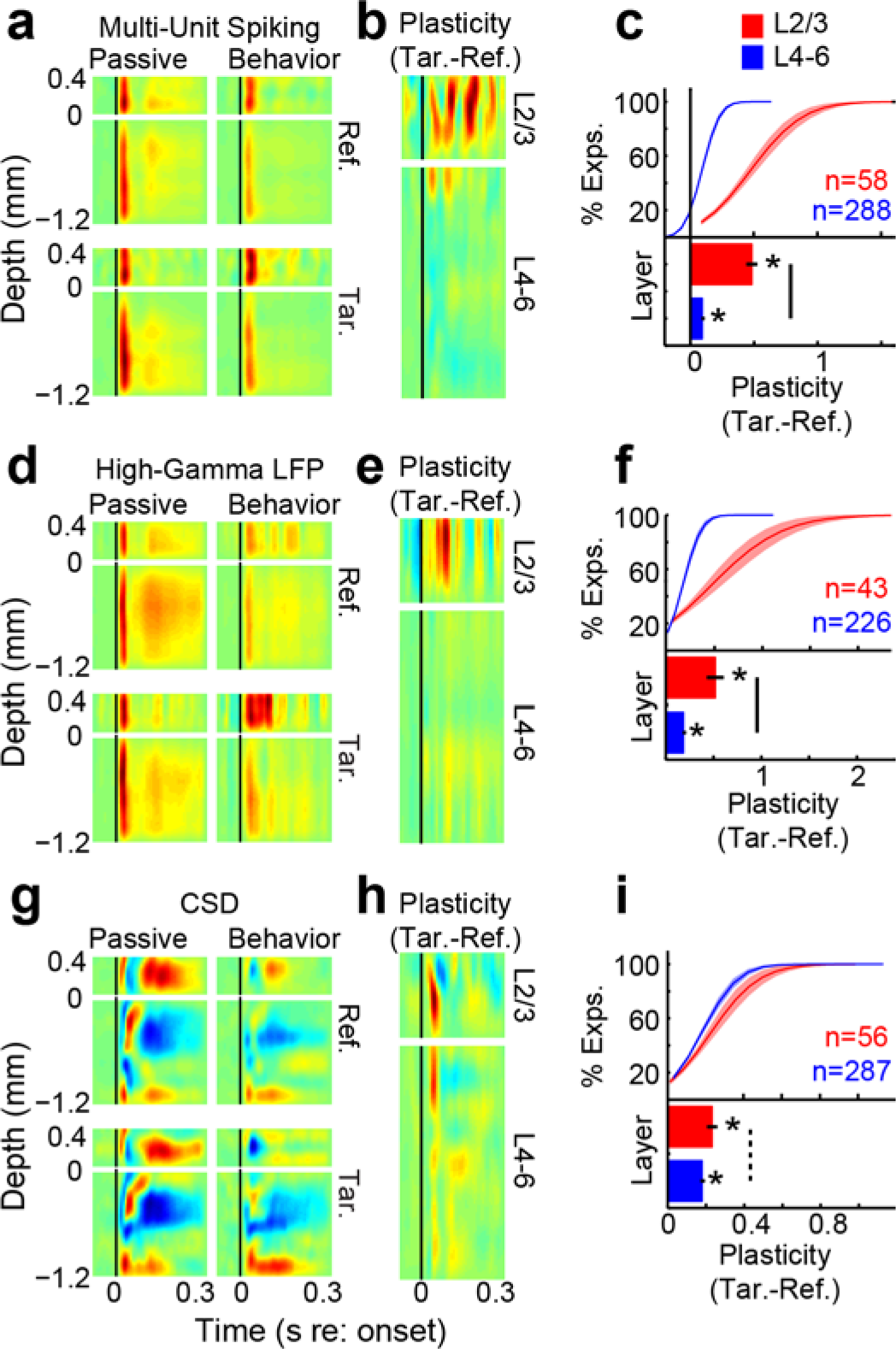
Laminar profiles of stimulus responses in primary auditory cortex (A1). The top, middle and bottom rows show the data for multi-units, high-gamma LFPs and current source densities (CSDs), respectively. Panels **a, d and g** show the average laminar profiles for reference noises (top row in each panel) and target tones (bottom row in each panel), and in passive (left column in each panel) and behavior (right column in each panel) conditions. Depth is marked relative to the L2/3 border (see Methods and Fig. 1c). For multi-units and high-gamma LFPs, red indicates an auditory response. For CSDs, red indicates a current source and blue indicates a current sink. Panels **b, e and h** show the laminar profile of rapid task-related plasticity, on the same depth axis as panels a, d and g. Red and blue indicates target enhancement and suppression, respectively, during task performance. Panels **c, f and i** show the cumulative distribution functions (top of each panel) and grand-averages (bottom of each panel) of task-related plasticity for all electrodes in L2/3 (red) and L4-6 (blue). Error bars and shading show 1 standard error of the mean (sem). Stars indicate averages that were significantly different than 0 (p<0.001, bootstrap t-test). Solid and dotted bars indicate significant differences between layers (p<0.001 and p<0.05, respectively, bootstrap t-test). Population sizes (n) indicate the number of electrodes per average after applying noise rejection (see Methods).

### Enhanced target selectivity in STRFs was greatest in L2/3

Task-related plasticity has previously been described in A1 using STRFs computed from single- and multi-unit spiking^4,6,7^. Here, we extend that analysis by computing STRFs from high-gamma LFPs (middle row, Fig. 3) and CSDs (bottom row, Fig. 3), in addition to multi-unit spiking (top row, Fig. 3). STRFs computed from reference noises estimate the magnitude of neural responses to target tones, relative to other pure-tone frequencies. We analyzed the 2-dimensional STRFs in the same manner as 1-dimensional response traces to compute laminar profiles of task-related plasticity (i.e., P_STRF_; Fig. 3b,e,h), with the additional step of first aligning each STRF to the target frequency bin before averaging. We found that enhanced target selectivity (i.e., peaks in P_STRF_; red in Fig. 3b,e,h) was the predominant effect in STRFs. Enhancement was greater in L2/3 than in L4-6 (Fig. 3c,f,i). The average STRF target enhancement for electrodes in L2/3 vs. L4-6 was: 0.6 vs. 0.27 (multi-unit spiking, p<0.001); 0.46 vs. 0.28 (high-gamma LFP, p<0.01); and 0.26 vs. 0.18 (CSD, p<0.01). Despite the different underlying neural mechanisms of multi-unit spiking, high-gamma LFPs, and CSDs^20–22^, the STRF plasticity that resulted in target enhancement was common to all three measures, i.e., reduced inhibitory fields (blue in Fig. 3a,d,g) and increased excitatory fields (red in Fig. 3a,d,g). This can be seen by comparing the left and right columns in panels a, d, and g in Figure 3. The STRF prediction of 27% target enhancement in multi-units from middle-deep L4-6 (Fig. 3c) is in agreement with previous measurements of task-related plasticity in A1 multi-unit STRFs^4,7^. Thus, we found that multi-unit spiking, high-gamma LFPs, and CSDs are similarly predictive of the effects of selective attention on A1 responses to behaviorally meaningful sounds.

**Figure 3.**
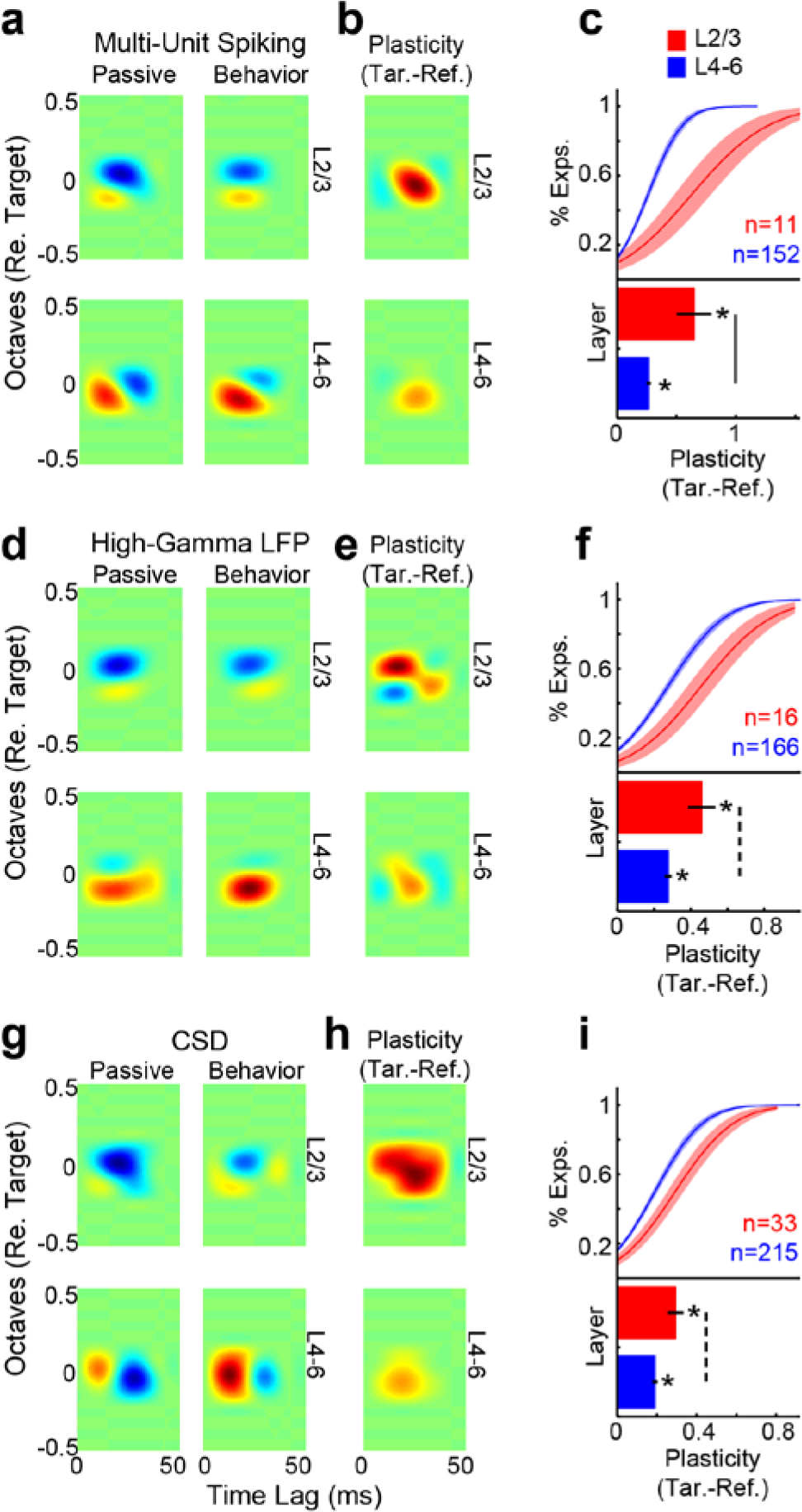
Laminar profiles of spectrotemporal receptive fields (STRFs) in A1. The top, middle, and bottom rows show the data for multi-units, high-gamma LFPs, and CSDs, respectively. Panels **a, d and g** show the average depth-registered STRF laminar profiles for L2/3 (top row in each panel) and L4-6 (bottom row in each panel), and in passive (left column in each panel) and behavior (right column in each panel) conditions. Red and blue indicate excitatory and inhibitory fields, respectively. Panels **b, e and h** show the laminar profile of task-related plasticity. Red and blue indicate target enhancement and suppression, respectively, during task performance. Panels **c, f and i** show the CDFs (top of each panel) and grand-average (bottom) of task-related plasticity for all electrodes in L2/3 (red) and L4-6 (blue). Error bars and shading show 1 sem. Stars indicate averages that were significantly different than 0 (p<0.001, bootstrap t-test). Solid and dashed bars indicate significant differences between layers (p<0.001 and p<0.01, respectively, bootstrap t-test). Population sizes (n) indicate the number of electrodes per average after applying noise rejection (see Methods).

### The persistence of target enhancement was common across cortical layers

Task-related plasticity can persist for minutes to hours after task performance ends^4,7^. We measured the persistence of task-related plasticity in the minutes following task performance, when the animal returned to a passive, quiescent state (i.e. during a “post-passive” condition). In figures 2 and 3, task-related plasticity was found, for both neural responses and STRFs, by computing Tar_Bhv/Pass_ - Ref_Bhv/Pass_. We quantified the persistence of task-related plasticity similarly by comparing the post-passive state vs. the “pre-passive” state that occurred before task performance, i.e., we computed P_persistence_ = Tar_Pre/Post_ - Ref_Pre/Post_. We found a similar pattern of persistence in both the neural responses (Fig. 4a) and STRFs (Fig. 4b) computed from multi-unit spiking, high-gamma LFPs, and CSDs: target enhancement was greatest during task performance and tended to decrease toward the pre-passive state after task performance.

**Figure 4.**
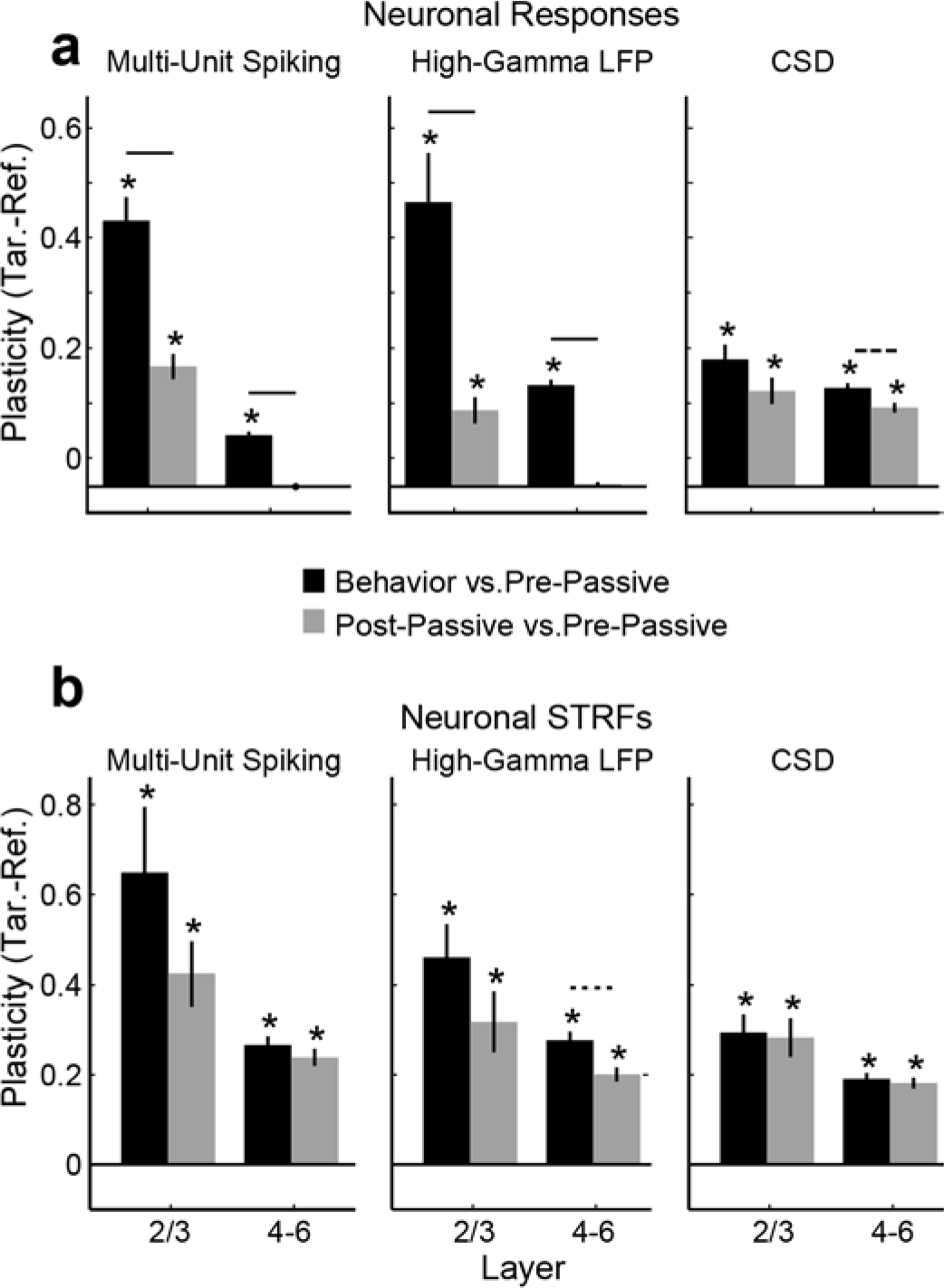
The persistence of task-related plasticity. Panels **a and b** show the persistence of task-related plasticity (i.e. target enhancement) for both neural responses and STRFs, respectively. The left, middle, and right columns of each panel show the results for multi-unit spiking, high-gamma LFPs, and CSDs, respectively. We found that target enhancement often persisted after task performance but was usually less than during the task. Stars indicate averages that were significantly different than 0 (p<0.001, bootstrap t-test). Error bars show 1 sem. Solid, dashed, and dotted lines indicate significant decay of task-related plasticity (p<0.001, p<0.01, p<0.05, respectively).

## Discussion

We recorded laminar profiles of neural activity in A1 during the performance of a pure-tone detection task and found that task-related plasticity was greater in L2/3 than in L4-6. The predominant effect of task-related plasticity was to enhance both neural responses to auditory targets and STRF selectivity for auditory target frequencies. We also found that target enhancement was similar for three different measures of neural activity: multi-unit spiking, high-gamma LFPs, and CSDs.

The dominance of target enhancement in L2/3 suggests that intracortical modulation of stimulus selectivity in A1 is an important neural correlate of selective attention. Top-down projections from prefrontal cortex are known to target neurons in supragranular layers in auditory cortex^24–26^. Neurons in prefrontal cortex show greater selectivity than A1 for behaviorally meaningful sounds^15^, and stimulation of orbitofrontal cortex causes changes in A1 pure-tone frequency tuning^16,27^that resembles the task-related plasticity observed here and in previous studies^4–7^. Simultaneous recordings from frontal cortex and auditory cortex reveal behavior-dependent changes in functional connectivity^13,15^.

Figure 2g shows that the greatest target enhancement measured from L2/3 CSDs arose from an increase in the magnitude of the target-evoked CSD current sink (blue values) between 40-100 ms. This long latency for maximal target enhancement in L2/3 also indicates the importance of intracortical, rather than thalamocortical, connections in task-related plasticity. Furthermore, target enhancement tended to be greater in multi-unit spiking and high-gamma LFPs than in CSDs (see Figs. 2-4). Since CSD current sinks represent input to the local network^20–22,28^, and spike metrics from multi-units and high-gamma LFPs represent local network output, it may be that intracortical input to A1 initiates or maintains target enhancement, which is then amplified by local neural populations. Future studies measuring task-related plasticity simultaneously in laminar profiles of A1 and higher-order cortex will help to clarify the intracortical network dynamics of target enhancement during auditory tasks.

Our study supports the growing body of evidence suggesting the importance of circuitry in L2/3 for plasticity in sensory cortex^29–32^. Our results also suggest that intracortical modulation of auditory processing is important not only for establishing long-lasting experience-related plasticity^19,33^but also for enabling rapid task-related plasticity as a neural correlate for selective attention.

## Methods

Neural activity was recorded in primary auditory cortex (A1) of 2 awake, behaving ferrets during 24 total experiments (12 experiments per animal). All experimental procedures were approved by the University of Maryland (UMD) Animal Care and Use Committee, and performed in accordance with UMD and National Institutes of Health guidelines and regulations.

Animals were trained to detect a pure-tone target after a series of references noises composed of temporally orthogonal ripple combinations (TORCs)^34^. Animals were initially trained in sound-attenuated testing booths where they could move freely. Once they reached behavioral criterion on the task (discrimination ratio > 0.6), they were implanted with a head-post and trained to perform the task while their heads were held fixed to facilitate stability in neurophysiological recording. Behavior and stimulus presentation were controlled by custom software written in Matlab (MathWorks).

### Acoustic stimuli

Target tones were pure sine waves (5-ms onset and offset ramps), with frequency held fixed during a block of trials, but varied randomly between experiments. Reference noises consisted of a set of TORCs with a spectral resolution of 0-1.2 cycles/octave and temporal envelope resolution of 4-48 Hz^34^. Targets and references always had the same duration (2 s, 0.8 s inter-stimulus interval) and sound level (65 to 80 dB SPL) during neurophysiological recordings. All sounds were synthesized using a 44 kHz sampling rate, and presented through a free-field speaker that was equalized to achieve a flat gain.

### Pure-tone detection task

Two animals were trained to perform a conditioned avoidance Go/No-go pure-tone detection task^35^(Fig. 1a,b). Training was initiated by delivering water from a spout while presenting reference noises. The animals quickly learned to freely lick the spout during references. Target tones were then introduced and the animals learned to stop licking the spout in a 0.4 s time-window after the target to avoid a mild shock to the tongue (free-moving behavior) or to the tail (head-fixed behavior). On each trial, the number of references presented before the target varied randomly from one to six. Catch trials were also used, in which targets were absent. Performance was assessed by the sensitivity index, d’, calculated from the probability of hits (reduced licking after target offset) vs. false alarms (reduced licking after reference offset)^36^.

### Neurophysiology

Each animal was implanted with a steel head-post to allow for stable recording, and a small craniotomy (1–2 mm diameter) was opened over A1. Recordings were verified as being in A1 according to their tonotopic organization, auditory response latency, and simple frequency tuning. Data acquisition was controlled using the MATLAB software MANTA^37^. Neural activity was recorded using a 24 channel Plexon U-Probe (electrode impedance: ~1 MOhm, 75 μm inter-electrode spacing). The probe was inserted through the dura, orthogonal to the brain’s surface, until the majority of channels displayed spontaneous spiking.

### Extracting neural responses

Multi-unit spikes were extracted on each electrode by band-pass filtering the raw signal between 300 and 6,000 Hz, then isolating spikes by peak detection (4σ threshold). Peri-stimulus time histograms (PSTHs) of spiking were computed using 10 ms bins. We analyzed multi-units instead of single-units because previous reports have indicated that task-related plasticity is more robust for multi-units^4^, which emphasizes that the behavioral relevance of task-related plasticity is predominant in neural populations, rather than single-units.

On the same electrodes used to extract multi-units, we also extracted high-gamma local field potentials (LFPs) by filtering the raw signal between 80-300 Hz, then taking the magnitude of the filtered signal’s Hilbert transform, and finally low-pass filtering below 70 Hz. Current source densities (CSDs) were derived by low-pass filtering the raw signal below 80 Hz, then computing the second derivative approximation of the filtered LFP across electrodes on the linear array^21^. LFPs on a given electrode were only kept if the signal-to-noise ratio (SNR) was greater than 1. This criterion eliminated 2 of 24 experiments from the dataset.

### Computing spectrotemporal receptive fields (STRFs)

STRFs were estimated by reverse correlation between each time-varying neural response (i.e., multi-unit spiking, high-gamma LFPs, and CSDs) and the TORCs presented during experiments^20^. Positive STRF values indicate time-frequency components of the TORC that correlated with increased neural responses (i.e., an excitatory field), and negative values indicate components that correlated with decreased responses (i.e. an inhibitory field). An STRF was only included in further analyses if its SNR was above the 25th percentile of the SNR distribution. Before averaging STRFs across electrodes, we aligned each STRF so that the frequency bin containing the target was in the center of the frequency axis.

### Depth-Registration

Each penetration of the linear electrode array produced a laminar profile of auditory responses in A1 across a 1.8 mm depth, however, the absolute depth varied across penetrations. In order to align all penetrations to the same depth, LFP responses to 100 ms tones were measured during the passive condition to find a common marker of depth (Fig. 1c). The marker was found for each penetration by first identifying the electrode with the shortest LFP response latency (Eτ), indicating an electrode depth at thalamorecpient layer 4. We then found the correlation coefficient between the average LFP waveform from Eτ and the LFP waveforms on all other electrodes in the same penetration. The border between the first neighboring electrode pair with positive and negative correlation coefficients defined the superficial vs. middle-deep layer border, corresponding to layer 2/3 (L2/3) and L4-6, respectively^12,22,23,38–40^. Laminar profiles were averaged across penetrations by aligning to the calculated border. Because of the neural response SNR criterion, data from the top two electrodes were also eliminated from all experiments, which removed data that may have included layer 1^23^. Thus, we were able to measure 1.6 mm laminar profiles that included layers 2/3-6^12,22,23,38–40^. We did not include 4 of the remaining 22 penetrations because the LFP correlations became negative in deep electrodes, suggesting that the penetration was not orthogonal to the surface or to the cortical layers.

### Quantifying the laminar profile of task-related plasticity

To quantify task-related plasticity (P) from neural responses (P_Resp_), we first computed the ratio of response amplitudes during behavior vs. passive trials, separately for target (Tar_Bhv/Pass_) and reference (Ref_Bhv/Pass_) sounds. Then we took the difference between target and reference ratio (P_Resp_ = Tar_Bhv/Pass_ - Ref_Bhv/Pass_). P_Resp_ was normalized between +/− 1 for each experiment before averaging across experiments. Positive values of P_Resp_ indicate target response enhancement during auditory task performance. We analyzed the 2-dimensional STRFs in the same manner as 1-dimensional response traces to compute STRF laminar profiles of task-related plasticity (i.e., P_STRF_), with the additional step of first aligning each STRF to the target frequency bin before averaging. Data from electrodes in each penetration were separated into either L2/3 or L4-6 STRFs, since these were the regions quantitatively defined by depth-registration. Significant differences between P_STRF_ and P_Resp_ from L2/3 vs. L4-6 were determined by a bootstrap t-test, using 100,000 resampling iterations. We estimated cumulative distribution functions (CDFs) for task-related plasticity by bootstrapping parametric fits of a Gaussian CDF to the data from each experiment.

### Data Availability

The dataset is available from the corresponding author on request.

## Acknowledgements

NAF, BE, JBF and SS supported by NIH R01DC009607 and R01DC005779. NAF supported by NIH/NIDCD F32DC013722 and T32DC00046. DE supported by CONICYT Becas Chile Doctorado (72100839), Postdoctorado (74170109), and Fulbright/IIE.

## Author Contributions

NAF, DE, BE, JBF and SS designed research. DE and BE performed experiments. NAF analyzed data and wrote the original draft of the manuscript. NAF, DE, JBF, and SS edited the manuscript.

